# Semi-automatic Geometrical Reconstruction and Analysis of Filopodia Dynamics in 4D Two-Photon Microscopy Images

**DOI:** 10.1101/2025.05.20.654789

**Authors:** Blaž Brence, Josephine Brummer, Vincent J. Dercksen, Mehmet Neset Özel, Abhishkek Kulkarni, Neele Wolterhoff, Steffen Prohaska, Peter Robin Hiesinger, Daniel Baum

**Author notes:** Corresponding authors. E-mails: Blaž Brence -; Daniel Baum. These authors contributed equally to this work (Blaž Brence, Josephine Brummer).

## Abstract

**Background:** Filopodia are thin and dynamic membrane protrusions that play a crucial role in cell migration, axon guidance, and other processes where cells explore and interact with their surroundings. Historically, filopodial dynamics have been studied in great detail in 2D in cultured cells, and more recently in 3D culture as well as living brains. However, there is a lack of efficient tools to trace and track filopodia in 4D images of complex brain cells.

**Results:** To address this issue, we have developed a semi-automatic workflow for tracing filopodia in 3D images and tracking the traced filopodia over time. The workflow was developed based on high-resolution data of photoreceptor axon terminals in the in vivo context of normal *Drosophila* brain development, but devised to be applicable to filopodia in any system, including at different temporal and spatial scales. In contrast to the pre-existing methods, our workflow relies solely on the original intensity images without the requirement for segmentation or complex preprocessing. The workflow was realized in C++ within the *Amira* software system and consists of two main parts, dataset pre-processing, and geometrical filopodia reconstruction, where each of the two parts comprises multiple steps. In this paper, we provide an extensive workflow description and demonstrate its versatility for two different axo-dendritic morphologies, R7 and Dm8 cells. Finally, we provide an analysis of the time requirements for user input and data processing.

**Conclusion:** To facilitate simple application within *Amira* or other frameworks, we share the source code, which is available at https://github.com/zibamira/filopodia-tool.

## 1 Background

Filopodia are thin, spike-like protrusions that dynamically extend and retract from the surface of cells and neuronal axon terminals. Filopodial dynamics are often interpreted as exploratory and may aid in directed growth, amongst other functions [1–3]. Neurons develop complicated morphologies through the slow path-finding of filopodia-rich axon terminal structures and branched dendritic structures that are preceded by continuously extending and retracting filopodia. While filopodial dynamics are thus intuitively associated with cellular morphogenesis and growth, the precise types of filopodial dynamics and their effects on biological functions has rarely been studied quantitatively based on live data that capture dynamics at sufficient resolution in time and space.

To address this issue, we have previously developed a *Drosophila melanogaster* brain culture live imaging system that takes advantage of the fly’s brain size, genetic tools, and limited time period of brain development [4]. Recently, fast filopodial dynamics have also been measured inside the intact developing *Drosophila* pupal brain [5]. In short, two-photon microscopy allows to visualize the in-vivo dynamics of filopodia anywhere in the *Drosophila* brain at high spatial and temporal resolution throughout the entire time window of axon pathfinding and synapse formation (for up to 2 days). Similar live observations of filopodial dynamics have been achieved in model systems ranging from 3D culture [6] to mice [7] and birds [8]. Advances in two-photon microscopy and newer imaging technologies like lattice-lightsheet microscopy increasingly allow to obtain such high-resolution data, leading to an analytical bottleneck.

To facilitate the quantitative analysis of *Drosophila* filopodial dynamics and their comparison across different experimental conditions, we developed computational approaches based on 3D time series of photoreceptor axon terminals (see Fig. 1). The workflow has been successfully applied for the analysis of filopodial dynamics in several studies [9–11].

**Fig. 1:**
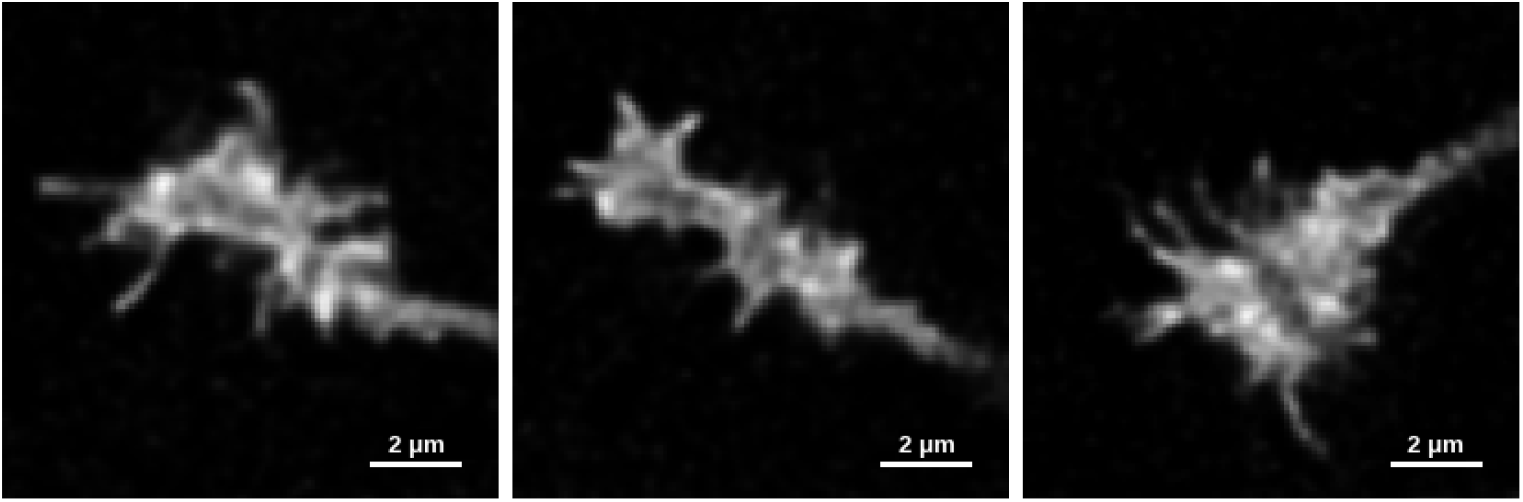
2D cross-sections of 3D two-photon microscopy images of different axon terminals with filopodia.

To analyze and compare filopodia dynamics in different datasets, an efficient method for information extraction and quantification is required. Of particular interest is information about filopodia length and lifetime as well as the number of extension or retraction events. To obtain such information, a 4D geometrical reconstruction of the filopodia geometry tracked over time is required. While the 4D images (3D + time) acquired using two-photon microscopy provide a high resolution in the first two dimensions (see Fig. 2a), the limited resolution in the third dimension constitutes a challenge (see Fig. 2b).

**Fig. 2:**
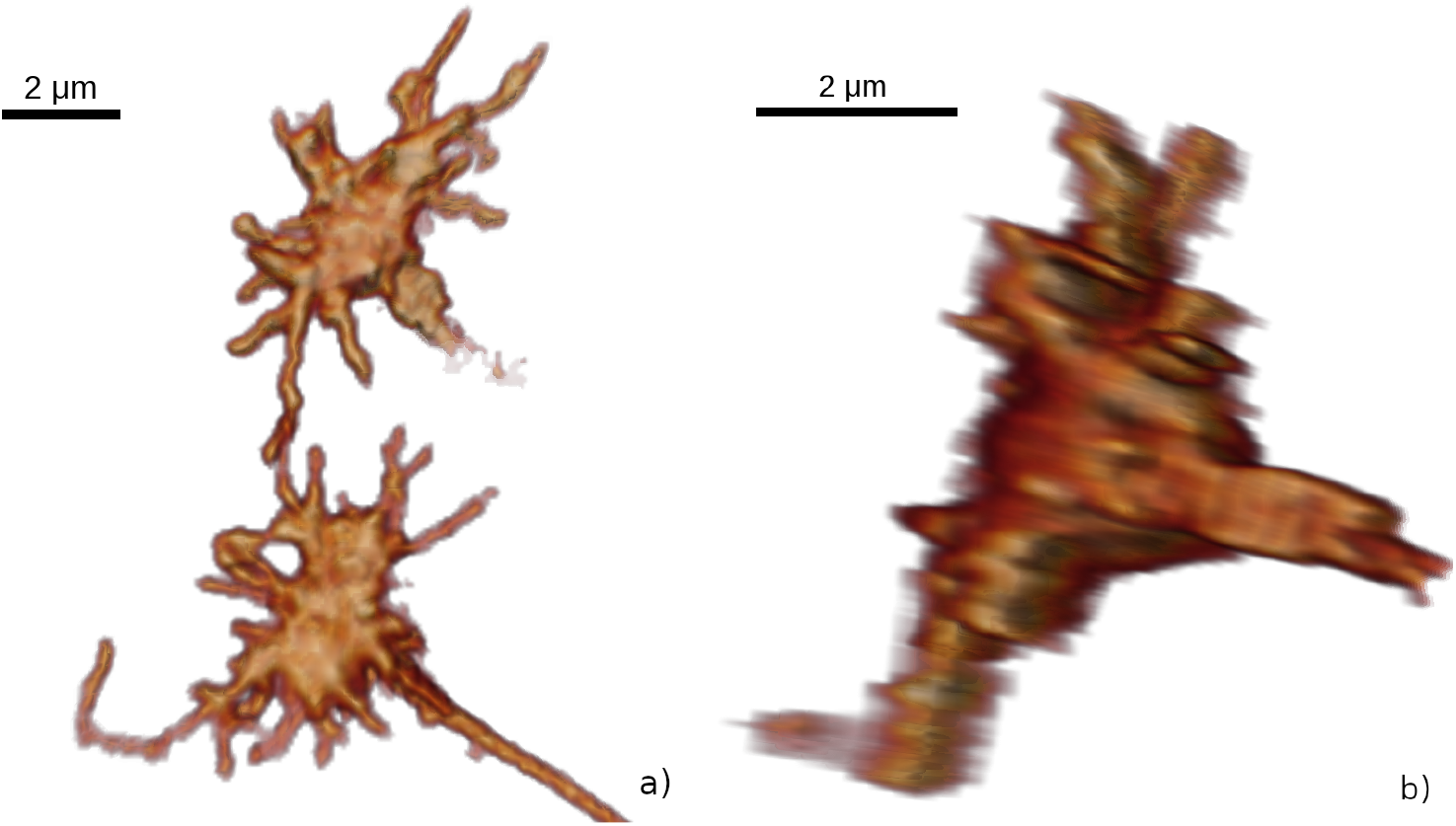
3D images of example axon terminals showing filopodia, visualized using volume rendering. a) Two-photon microscopy images of two axon terminals shown in x-y orientation. b) The axial resolution in z direction (here the horizontal direction) is poorer (0.1 × 0.1 × 0.5 *µm*^3^).

There are several existing software solutions to analyze filopodia. These can be divided into general purpose protrusion tracing and tracking tools, and tools designed specifically for the study of filopodia. General tools are mostly dedicated to determining the cell shape [12] or general cellular protrusion analysis [13, 14]. As filopodia are small, high-resolution images are needed to ensure tracking efficiency with these tools [13]. Filopodia-specific tools typically offer similar analysis in terms of filopodia quantification. Previous studies demonstrated analyses of parameters including length, shape, density, elongation and retraction speed [15–21]. A few solutions additionally provide tools for spatio-temporal analyses of protein concentrations in filopodia [15, 18]. Importantly, all the tools described so far are based on 2D data and time, or 3D data with reduced dimensionality (working with 2D projections and time). To accurately analyze filopodial dynamics, 3D analysis must be performed. 3D tracing of filopodia can be achieved using one of the commercial microscopy image analysis tools, such as *Imaris*. Its *Filament tracer* module can be used to trace filopodia as well [4]. *TeraVR* is another tool that is capable of 3D geometrical neuron reconstruction [22]. It allows the user to work within immersive virtual reality, where one can visualize and interact with images. However, both *Imaris* and *TeraVR* do not allow for automatic filopodia propagation or automatic labeling of the same filopodium in successive time steps. At the time of this study, 3D analysis of filopodia or cellular protrusions over time was conducted only in a few publications [23–25]. These tools use convolutional neural networks for image segmentation, hence they need a potentially large amount of training data of diverse datasets to generalize for different types of data.

Here, we present a semi-automatic workflow for filopodia tracing and tracking that does not require training data or image segmentation. Instead, our workflow works directly with the intensity images as input. The workflow was developed within the 3D visualization and data analysis software *Amira* [26], extending its *Filament Editor* [27], but it can be implemented in other environments. The semi-automatic tool supports interaction in both 2D and 3D viewers, thus allowing for more intuitive and faster data processing. The results of the 4D geometrical filopodia reconstruction (referred to as ‘reconstruction’ throughout the paper for simplicity) are represented as skeleton graphs. Such graphs simplify the extraction of statistical information, such as filopodial length, number, orientation, retraction, and extensions. Using this semi-automatic approach, manual input is significantly reduced. We provide an open-source extension package, which allows implementation of this workflow in *Amira* and can serve as a guide for implementation in an alternative environment.

## 2 Implementation

The semi-automatic workflow and a dedicated set of tools have been developed within the *Filament Editor* framework [27] of *Amira*, as it already contains a lot of functionality necessary for its implementation. For example, it provides general 2D/3D visualizations (Fig. 3) and interactive editing of filamentous, graph-like objects embedded in 3D space. The 2D view displays an image cross-section with the graph superimposed and offers tools for interactive tracing. In addition, the 3D view can display the graph and a 3D direct volume rendering visualization of the image data. To support the tracing and tracking of filopodia over time, we extended the capabilities of *Amira*’s *Filament Editor* to process time-dependent geometry and support the specific functionality for the tracing of filopodia described below.

**Fig. 3:**
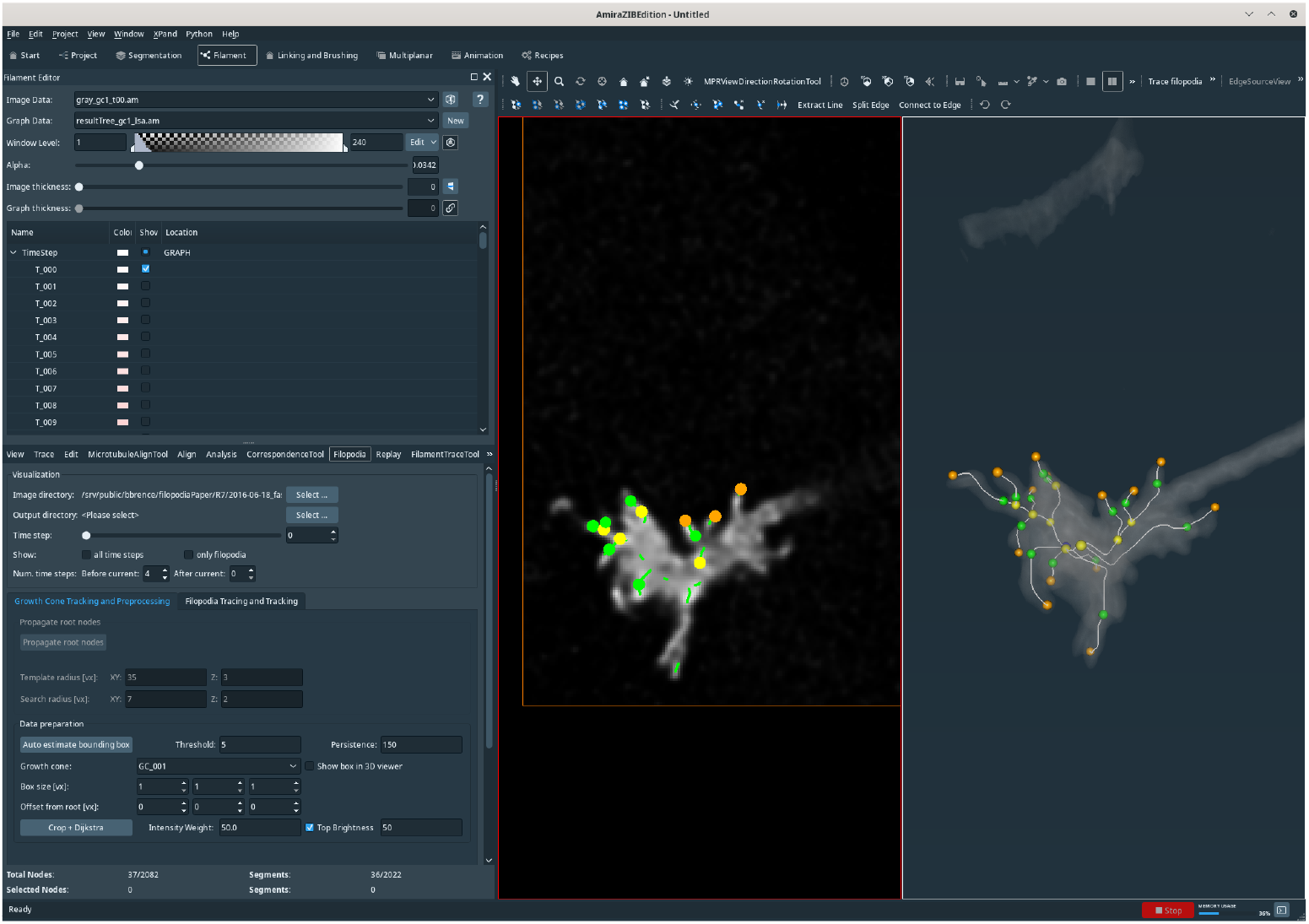
*Filament Editor* with 2D and 3D viewer. The *Filament Editor* provides a 2D (center) and 3D (right) viewer to display the neuronal axon terminals. Both 2D and 3D viewer show the skeleton graph superimposed in the image. Various tools allow the display, processing, and analysis of the filopodia.

### 2.1 Data acquisition

We developed the workflow based on live imaging data of R7 photoreceptor neurons and subsequently tested the methods for other neurons inside the intact *Drosophila* brain. The datasets were imaged using two-photon microscopy, yielding 4D data (3D + time) with spatiotemporal resolution detailed enough to discern individual filopodial dynamics [4, 5]. Specifically, all datasets used here cover a period of 60 minutes with a temporal resolution of 1 minute. However, the images have a limited axial resolution in z direction that leads to an anisotropic voxel size of 0.1 × 0.1 × 0.5 *µm*^3^. Images were acquired with a Leica SP8 OPO-MPX multiphoton microscope using a 40x oil immersion lens.

### 2.2 Data representation

In order to facilitate tracking of filopodia and subsequent analysis, R7 photoreceptor axon terminals (including filopodia) are represented by their skeleton graphs (trees). Nodes of the graph have one of the following types, as shown in Figure 4:

**Fig. 4:**
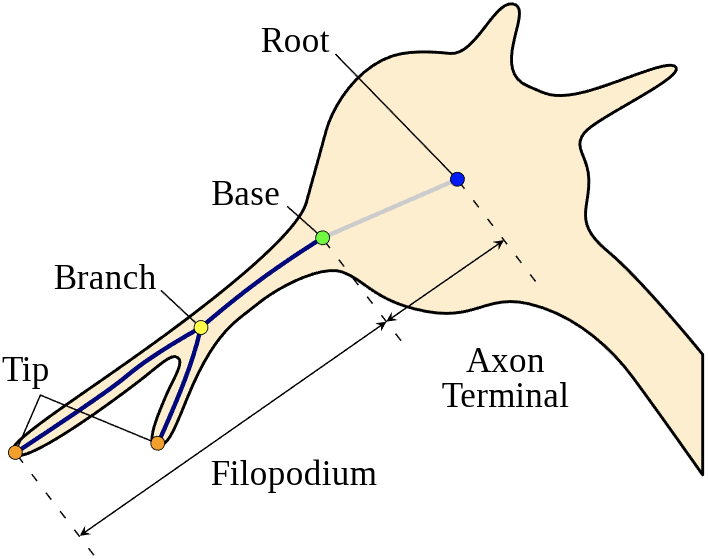
Schematic of the 3D tree-like structure of neuronal axon terminals. A neuronal axon terminal with its filopodial protuberances can be represented as a skeleton graph (tree). Here, a single branch of the tree is shown. Branches extend from the *root* node (blue) to the filopodia *tips* (orange), passing through the *base* locations (green) and, potentially, *branching* nodes (yellow), that is, when these exist. The part of the path between tip and base is the *filopodium*; the part of the path between base and center is labeled as *axon terminal*.

1. Root node (axon terminal center)
2. Base node
3. Branching node
4. Tip node

Root, base and tip nodes are mandatory when tracing filopodia whereas branching nodes are not because filopodia branching happens rarely.

Edges of the graph connect the nodes. Edges that are part of the paths between root and base are labeled as *axon terminal*, the ones between base and tip as *filopodium*. The skeleton graphs of all time steps are combined in one dataset. All nodes and edges obtain a time label to mark the time step of their occurrence.

### 2.3 Workflow

The overall workflow consists of the following steps:

1. Preprocessing
  a. Determination of root nodes
  b. Extraction of axon images
2. Filopodia reconstruction (per axon)
  a. Filopodia tracing
  b. Filopodia tracking
  c. Proofreading
3. Statistical analysis

The individual workflow steps are introduced and explained in the following sub-sections. All user-adjustable workflow parameters including parameter domain and parameter values used in this paper are listed in Table 1.

**Table 1:** User-adjustable parameters. Overview of the user-adjustable parameters used during the workflow execution. The images of the used datasets have integer intensity values in the range [0, 255]. The template and search window sizes *T* and *S* for the node propagation depend on the cropped image size, which can be different for each axon terminal. *θ, p* and *I*_*max*_ depend on the range of intensity values in the images. Threshold distance *d* is the maximum distance that two nodes can be apart from each other to be considered matching and can be connected in the tracking process.

| Task | Acr. | Parameter | Domain | Used value |
| --- | --- | --- | --- | --- |
| Preprocessing | $\theta$ | Threshold of CTS | $[0, 255] \subset \mathbb{N}$ | 50 |
| | $p$ | Persistence of CTS | $[0, 255] \subset \mathbb{N}$ | 150 |
| | $T_R$ | Root template size [vx] | $\mathbb{N}^3$ | $15 \times 15 \times 3$ |
| | $S_R$ | Root search window [vx] | $\mathbb{N}^3$ | $70 \times 70 \times 7$ |
| | $\gamma_R$ | Root NCC value | $[0, 1] \subset \mathbb{R}$ | 0.8 |
| Tracing | $c$ | Intensity weight | $\mathbb{R}$ | 50.0 |
| | $I_{max}$ | Max. brightness threshold | $[0, 255] \subset \mathbb{N}$ | 100 |
| Tracking | $d$ | Threshold distance [$\mu m$] | $\mathbb{R}$ | 1.0 |
| | $l$ | Min. filopodia length [$\mu m$] | $\mathbb{R}$ | 0.5 |
| | $T_B$ | Base template size [vx] | $\mathbb{N}^3$ | $50 \times 50 \times 10$ |
| | $S_B$ | Base search window [vx] | $\mathbb{N}^3$ | $10 \times 10 \times 4$ |
| | $\gamma_R$ | Base NCC value | $[0, 1] \subset \mathbb{R}$ | 0.8 |
| | $T_T$ | Tip template size [vx] | $\mathbb{N}^3$ | $10 \times 10 \times 4$ |
| | $S_T$ | Tip search window [vx] | $\mathbb{N}^3$ | $15 \times 15 \times 4$ |
| | $\gamma_T$ | Tip NCC value | $[0, 1] \subset \mathbb{R}$ | 0.8 |

#### 2.3.1 Preprocessing

Some preprocessing is required to facilitate the filopodia reconstruction in step 2. This preprocessing separates the axon terminals from another. It is done in two steps, as described below.

##### Determination of axon terminal centers

First, the user imports the image data into *Amira* (Fig. 5a). Within the 2D viewer, the user selects the center of an axon terminal to be processed in the first time step. Centers for the remaining time steps are automatically detected with a template matching algorithm [28]. For this, the image region around the root node in the previous time step is used as template. The root node is iteratively repositioned to all the voxel centers inside a user-defined search window. At each position, the similarity of the image with the template is computed using the normalized cross correlation (NCC), utilizing the Insight Toolkit (ITK) [29] implementation. NCC is used to account for possible intensity shifts. The algorithm automatically selects the voxel with the maximal NCC value as the new root node. If the NCC value is less than the chosen threshold *γ* ∈ [0, 1], the new location is considered unreliable and the location of the current time step is used. In such a case, the user is notified and it is advisable to carefully proofread the positions of base nodes. The template size *T*_*R*_ ∈ ℝ^3^ approximates the size of the axon terminal; the search window size *S*_*R*_ ∈ ℝ^3^ is set to the maximal expected drift of the root node (during the workflow, no other drift-correction steps are necessary). Both template size and window size are user-adjustable with respect to the anisotropic voxel size. Center nodes are assigned an axon terminal ID and a time step label. In the next step, the user verifies the center locations for all the time steps by displaying the nodes together inside a 3D semi-transparent volume rendering of the image (Fig. 5b).

**Fig. 5:**
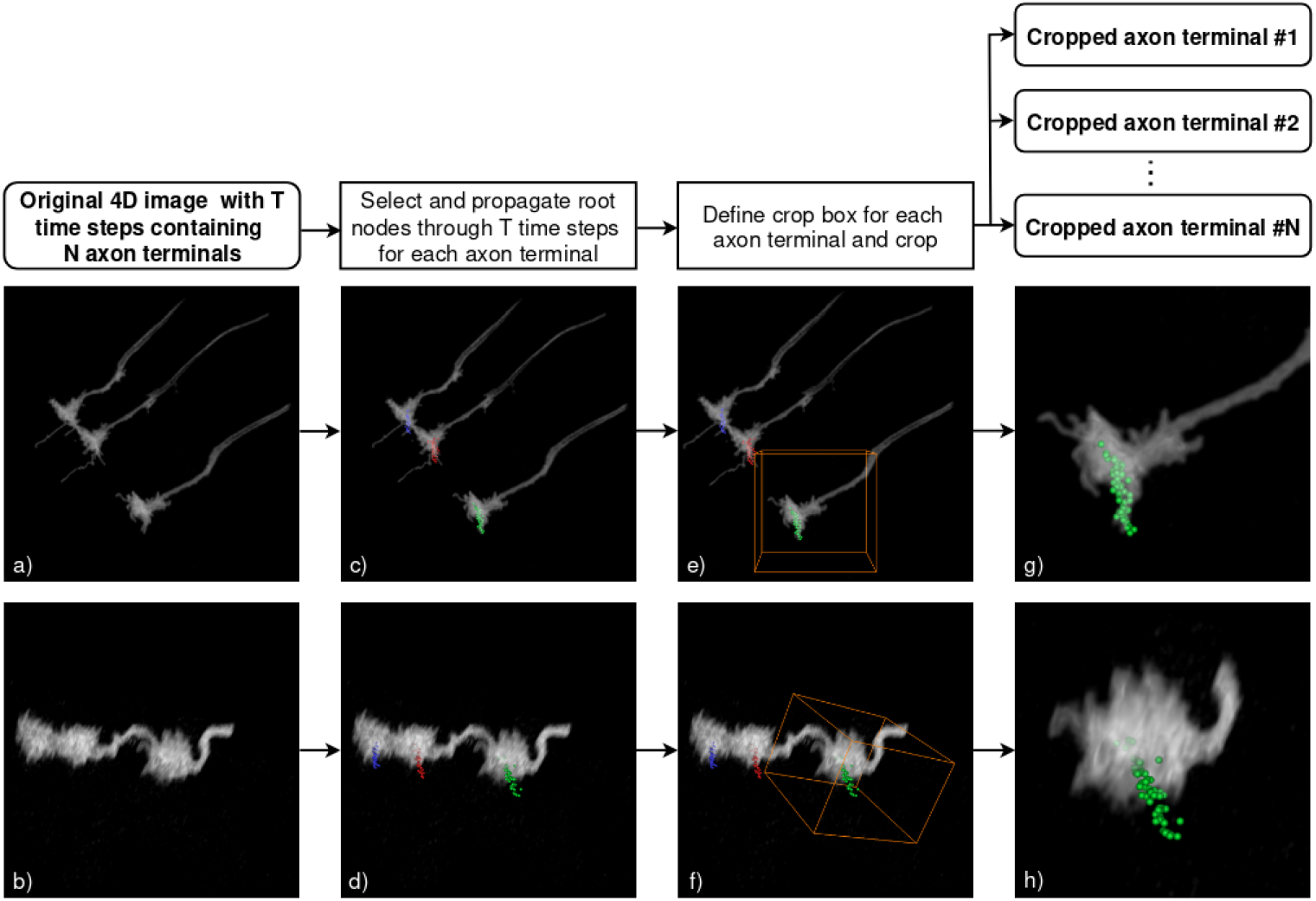
Preprocessing workflow. a, b) Images are loaded into *Amira*. c, d) User selects centers of the axon terminals to be analyzed. Centers are automatically propagated to the succeeding time steps. e, f) Crop box must be defined around each axon terminal. An initial cropping size is approximated with the Contour Tree Segmentation module and is user-adjustable afterwards. It is set to the minimal box including all voxels of the Axon terminal. g, h) Result of preprocessing are multiple datasets, each containing one axon terminal with marked centers and precomputed Dijkstra graphs. Upper and lower row show the same data but from different perspectives.

##### Extraction of axon terminal images

Filopodia are traced and tracked for one axon at a time. To facilitate this, cropped datasets are created containing single axons. Using cropped images significantly reduces computation time and reduces occlusion by other axons. The user can determine the size and location of the cropping box either numerically or by dragging the handles of a box in the 3D viewer. The same box size is used for all time steps. The origin of the box is shifted such that it always has the same position relative to the root node position. Initial estimation of the box location and its size are computed automatically. The estimation is based on the Contour Tree Segmentation (CTS) [30], using two user-adjustable parameters, intensity threshold *θ* and a persistence value *p*. Connected-component labeling of the binary image is not applicable since multiple axons can be very close to each other or even touch and, therefore, might not be separable using connected-component labeling.

The cropping box for a particular time step and axon is displayed in 3D, together with the root nodes and a volume rendering of the 3D image, allowing users to quickly verify whether the box completely surrounds the axon terminal (Fig. 5c). After visual validation, the images are cropped and saved (Fig. 5d). In addition, Dijkstra graphs [31] are pre-computed for each cropped image during the last step. They are used in the following steps.

#### 2.3.2 Filopodia reconstruction

The user traces the filopodia for one axon terminal at a time. The cropped images and pre-computed Dijkstra graphs of all time steps of a previously pre-processed dataset are loaded into the software. The process of filopodia reconstruction involves the following steps:

1. Filopodia tracing in one time step: A new filopodium is added by interactively specifying the position of its tip. The software determines the filopodium base automatically and traces the path from the tip node to the root node automatically.
2. Filopodia tracking across time steps: All filopodia, both new and previously propagated ones, are automatically propagated to the next time step.
3. Proofreading: The traced paths and tip locations, including the filopodia bases, are visually verified and interactively corrected if necessary.

All steps must be repeated as many times as necessary to reconstruct all filopodia in the desired amount of time steps. The full reconstruction workflow can be seen in Figure 6. The part of the workflow enframed with the red dotted line is described in more detail in Figure 7. The following sections contain detailed description of each reconstruction step.

**Fig. 6:**
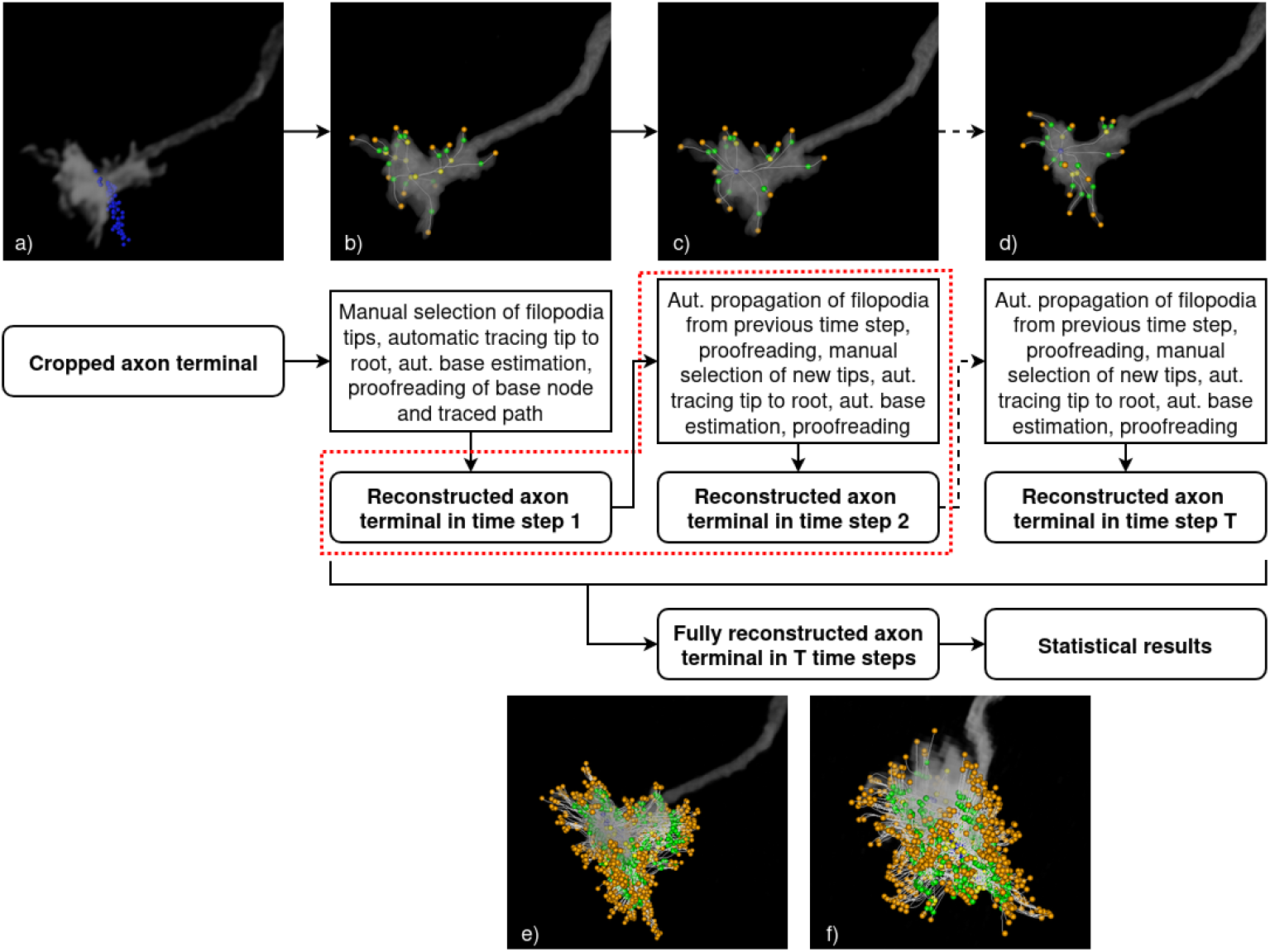
Reconstruction workflow. a) Reconstruction begins with loading dataset of a single cropped axon terminal and its graph, which at this point only contains the root nodes. b) In the next step, filopodia tips are manually selected and each filopodium is automatically traced to the root node. Traces and estimated base nodes must be proofread. c) Filopodia from time step 1 are then propagated to time step 2. All propagated filopodia must be proofread before selecting newly emerged tips and an automatic tracing to the root node. d) Process must be repeated for all time steps that are wished to be included in the analysis. e, f) From fully reconstructed axon terminal in all the time steps, statistical results are derived. Images show the same result from two different perspectives. The part of workflow enframed by the red dotted line is described in detail in Fig. 7.

**Fig. 7:**
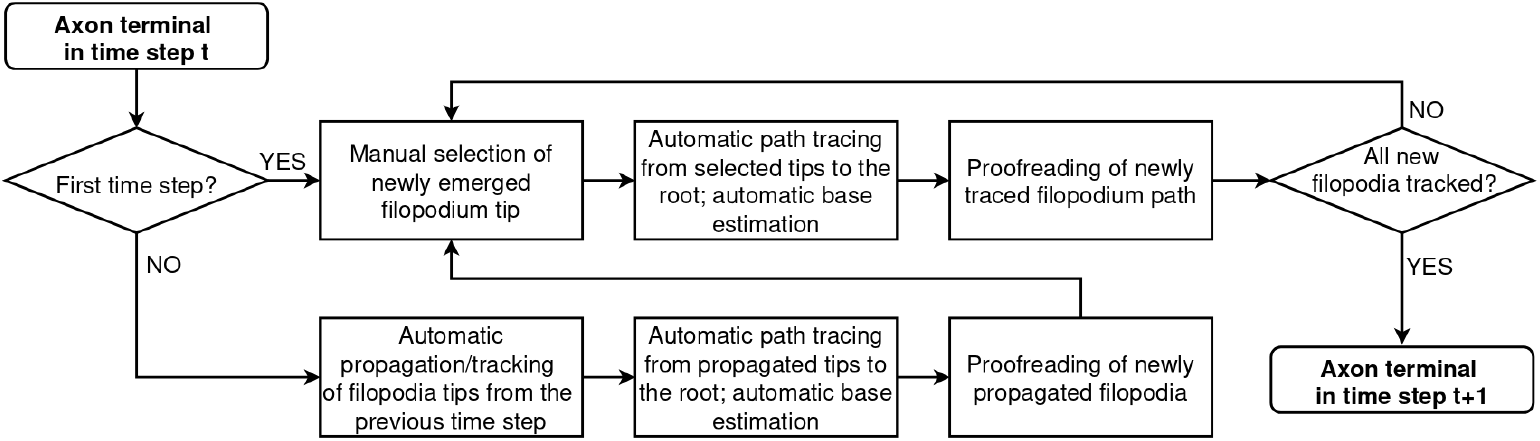
Reconstruction workflow. Generalized detailed workflow depicting reconstruction of an axon terminal between two consecutive time steps.

##### Filopodia tracing - path generation

The path between two nodes *N*_*B*_ and *N*_*T*_ is traced using an intensity-weighted Dijkstra shortest path algorithm [31]. A Dijkstra graph consists of nodes and weighted edges. Here, the set of nodes is given by the voxels contained in an axis-aligned box. This box is bounded by *N*_*B*_ and *N*_*T*_, extended in each dimension by *n* voxels to account for potential shortest paths that are not contained within original box bounded by *N*_*B*_ and *N*_*T*_. Empirically, we determined *n* = 10 to be a suitable value for our implementation. The set of edges consists of all connections between neighboring voxels (using the 26-neighborhood). The edge weight *w*_*ij*_ between two nodes *v*_*i*_ and *v*_*j*_ having intensity values *I*_*i*_ and *I*_*j*_ ∈ [0, 255] is defined as follows:

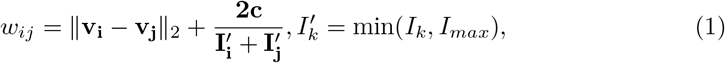

where

- **v**_**i**_, **v**_**j**_ are node coordinates,
- 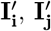 are the intensity values of the voxels **v**_**i**_ and **v**_**j**_, respectively, and
- **c** is a user-adjustable intensity weight.

As can be seen in Equation (1), the weights *w*_*ij*_ are computed using a distance term between the nodes and an intensity term penalizing dark voxels. The user-adjustable intensity weight **c** balances these terms. Empirically, **c** = 50 has been found to be a suitable value for the sample data used during development. To avoid that long paths through bright voxels are preferred over slightly shorter paths with darker foreground voxels, the foreground intensity is capped at *I*_*max*_, which is a user-adjustable parameter. Based on the tracing with many membrane-labeled filopodia in multiple datasets of various quality, we can assess that tracing from filopodial tips to root nodes is aided by brighter voxels along the path but also works robustly along paths that are only weakly labeled and have discontinuous voxel brightness.

To facilitate an efficient tracing from a point to the root node, Dijkstra graphs storing the shortest path tree (rooted at the axon terminal center) are pre-computed. The tree is encoded by storing the neighbour of each voxel that is next on the shortest path to the root. The shortest path from any voxel to the root can then be efficiently found by iteratively moving to the neighbor closest to the root until the root has been reached. To compute the path between a pair of non-root nodes, the Dijkstra graph is generated on the fly. This might be the case during graph corrections, for example.

To avoid coinciding paths, the algorithm performs an intersection test with the existing edges of the graph when adding a new path. Should an intersection be detected, a branching node is created. In case the branching node lies between the tip and the base, the branch integrates into the existing filopodium. If the filopodium branches between base and root node, it is treated as an independent filopodium.

##### Filopodia tracing - base estimation

After tracing a filopodium from a tip to the root node, the location of the filopodium base is automatically determined. Here, the assumption is made that the 2D intensity profile in a plane orthogonal to the traced path has Gaussian shape for the filopodium part and is non-Gaussian (ideally uniform) inside the axon terminal body. The base location is specified as the point where the intensity profile changes from Gaussian to non-Gaussian (Fig. 8c). The “Gaussian-ness” is estimated by computing the Root Mean Squared Deviation (RMSD) of radially sampled intensity values in a 2D plane orthogonal to the edge. Ideal Gaussian parameters are estimated from the sampled intensities. The RMSD error remains relatively constant inside a filopodium but increases drastically when entering the axon body (Fig. 8c).

**Fig. 8:**
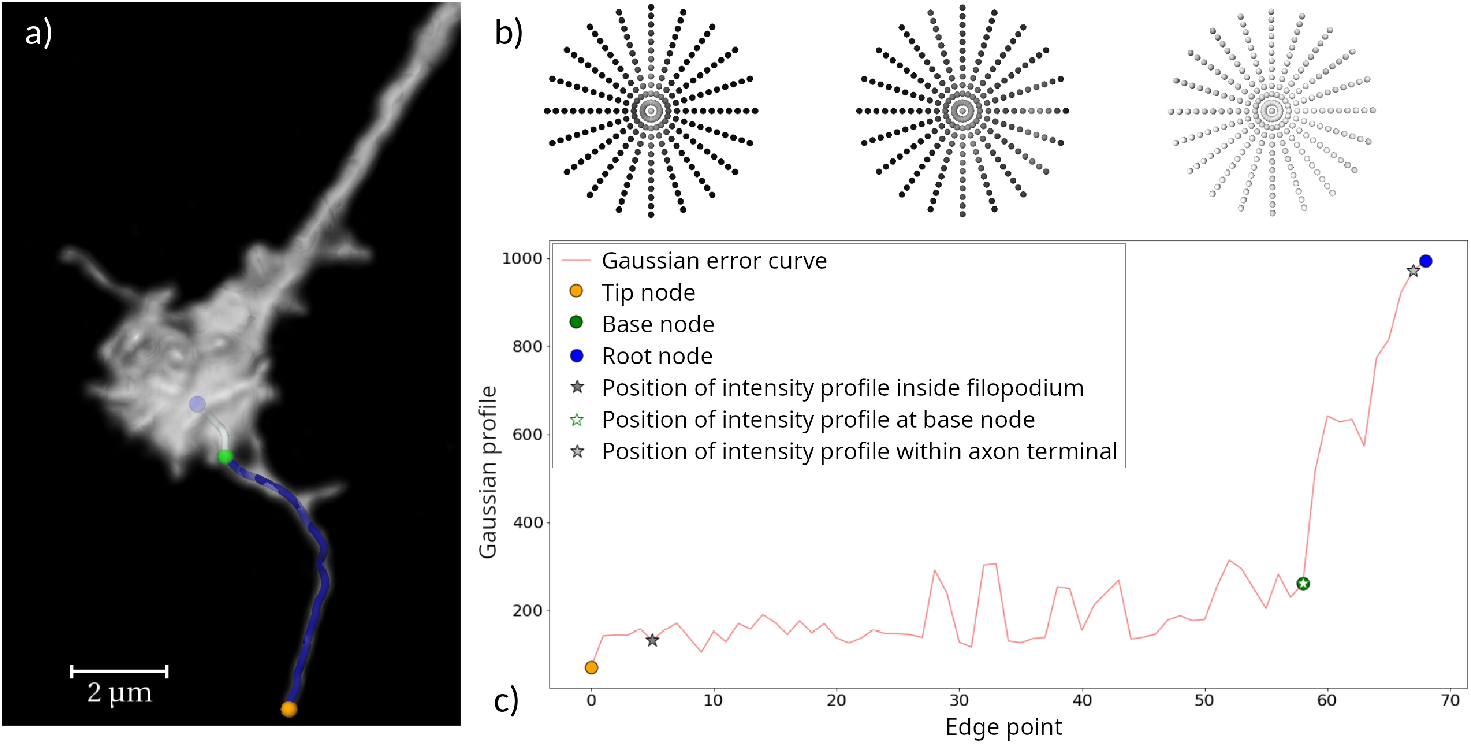
Adding a filopodium. a) The user adds a new filopodium by interactively specifying the tip (orange). The path to the root node (blue) is automatically traced. b) The base of the filopodium is detected by computing the Gaussian-ness of the intensity profile at each path point. From left to right: intensity profile from example point within filopodium, at base node (green), and from example point within axon terminal. c) The change from Gaussian-ness to non-Gaussian-ness indicates the base. The local peaks are caused by branches and the limited axial resolution in z.

##### Filopodia tracking

Filopodia are tracked over all time steps by automatic propagation. The propagation algorithm aims to detect the same filopodium in the successive time step. Filopodia tip and base are propagated using the same template matching algorithm used for propagating root nodes. Since root and base nodes rarely move, the search window for both node types can be small. In contrast, the filopodia tips show high motility. Therefore, the search window has to be adequately large to capture this movement. Template sizes differ as well since the high-intensity region around a tip will in general be much smaller than the high-intensity region around the base. Both search window and template size have to be determined experimentally and can be adjusted in the software. The paths between the nodes are traced as explained above in the *Filopodia tracing* section. The newly traced filopodium is assigned the same track ID as its “predecessor” in the previous time step.

In case a tip cannot be propagated (i.e. there is no location with NCC value above the threshold *γ*), the filopodium is considered to have retracted. If no base can be found, the location of the base in the previous time step is used. If the length of the path between base and tip is shorter than a user-specified threshold *λ*, the filopodium is also considered to have retracted, and is removed.

After propagation, the user verifies whether the automatically created traces are correct (see next paragraph). In addition, newly emerging filopodia are interactively added as before. These steps must be repeated until all time steps have been processed.

##### Proofreading

The automated tracing and tracking are not always accurate and especially minor corrections are common. Hence, interactive verification and correction is an important part of the workflow and is especially facilitated by the software tool. The following parameters need to be validated when moving through successive time steps:

- Location of tip and base
- Location of the traced path
- Location of any intersection points found
- Assignment of track ID

Dislocated base or tip nodes can be manually moved to new positions by selecting the node and clicking at the new location; incident edges are retraced automatically. Erroneous paths can be redirected by defining an additional supporting point or by selecting a point on an existing path; a branching node is then created at the intersection location. Incorrect track IDs can be corrected by explicitly selecting filopodia in subsequent time steps and marking them as matched. This is only the case when multiple filopodia are close to each other.

##### Workflow realization using low-level basic operations

The high-level tasks described above are realized by several basic operations that are implemented in the software:

- *add filopodium*: This operation is performed by defining a filopodia tip either in the 2D viewer (in either an orthogonal or an oblique slice) or in the 3D viewer using the volume rendering. The path is then automatically traced from the filopodium tip to the axon terminal center and the base is also automatically determined.
- *delete filopodium*: This operation is performed when a filopodium has been inaccurately propagated.
- *move tip*: Similar to the *add filopodium* operation, this operation can be done in the 2D or the 3D viewer.
- *move base*: This operation can be done in the 2D viewer or in the 3D viewer. The base can be moved to a completely new location or dragged along the existing edge.
- *move edge*: This operation is performed when the position of the traced path is found to be incorrect. In this case, further points can be added in 2D or 3D through which the path has to pass. The path is then retraced between the added points and the tip and base points.
- *propagate filopodia*: This operation is performed to propagate all filopodia from the current time step to the next. The operation is required after finishing time step reconstruction and is rarely invoked otherwise.
- *match filopodium*: This operation should be performed after adding a filopodium that was mistakenly not automatically propagated. The operation ensures that the same filopodium at consecutive time points has the same ID. It can also be used if filopodia in consecutive time steps are wrongly matched.

#### 2.3.3 Statistical analysis

The final result of the geometric reconstruction is a 4D skeleton graph representing the axon terminal and all filopodia for each time step. The following quantities are computed from the graph:

- Length of each filopodium in each time step
- Number of extension and retraction events
- Mean extension and retraction velocities
- Mean length and lifetime for each filopodium (through different time steps)
- Filopodia growth angles

The length is calculated as the sum of the Euclidean distances between consecutive edge points on the traced path. In case a filopodium branches between tip and base, the length of the branch is added to the filopodium length. From changes in length over time, we compute the number of extensions (filopodium is longer than in the previous time step) and retractions (filopodium is shorter than in the previous time step). The filopodium angle is defined as the angle between the unit vector and a vector spanning from the root to the base node at the time of their initial formation. The angle is computed as:

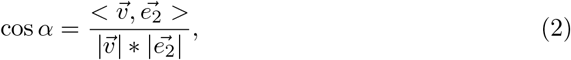

where 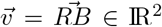 is the vector spanned from the root node *N*_*R*_ (axon terminal center) to the base node *N*_*B*_ of the filopodium, both projected onto the x-y plane and 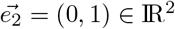 is the unit vector.

Since the coordinates of all nodes (root, base, tip) are known, the calculation of other angles can easily be performed. The mentioned filopodia growth angle is of interest as it provides information about the distribution of filopodia (bases) around the axon terminal. All computed quantities are stored in spreadsheets, which allow further validations and analyses with other tools, such as *Python, MATLAB* or *R*.

## 3. Results

The semi-automatic workflow was devised to fulfill two main requirements: first, to facilitate efficient 3D reconstruction of neuronal axon terminals and their filopodia; second, applicability to a wide range of datasets. To benchmark the efficiency of the method, we analyzed the time necessary for dataset preprocessing. To test applicability, we tested the method on a neuronal cell type with very different and much more elaborate axo-dendritic morphology.

### Analysis of preprocessing

As described above, the necessary preprocessing includes the determination of root nodes, image cropping, and the computation of the Dijkstra graphs. Table 2 shows the time taken for different steps of the preprocessing for a single user. In the data used for tool and workflow development, the entire preprocessing only took 3-4 minutes per dataset, while the actual filopodia tracing and tracking can be on the scale of hours.

**Table 2:**
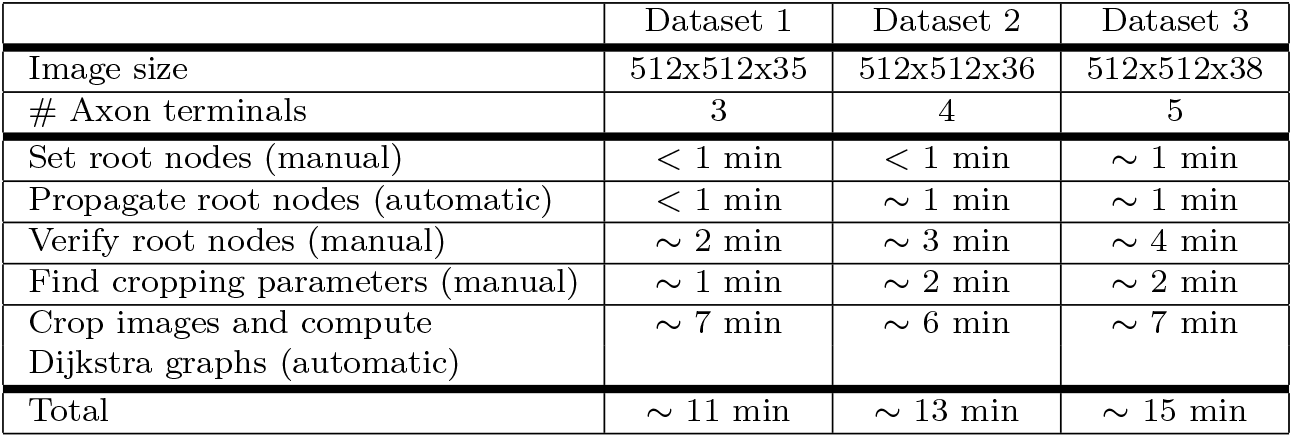
Time requirements for preprocessing steps for R7 cells. Preprocessing of a dataset normally depends on its size and number of axon terminals, and in the case of data analyzed here, it took less than 15 minutes per sample. Different steps are distinguished between automatic ones and those that are performed manually.

|  | Dataset 1 | Dataset 2 | Dataset 3 |
| --- | --- | --- | --- |
| Image size | 512x512x35 | 512x512x36 | 512x512x38 |
| # Axon terminals | 3 | 4 | 5 |
| Set root nodes (manual) | < 1 min | < 1 min | ~ 1 min |
| Propagate root nodes (automatic) | < 1 min | ~ 1 min | ~ 1 min |
| Verify root nodes (manual) | ~ 2 min | ~ 3 min | ~ 4 min |
| Find cropping parameters (manual) | ~ 1 min | ~ 2 min | ~ 2 min |
| Crop images and compute Dijkstra graphs (automatic) | ~ 7 min | ~ 6 min | ~ 7 min |
| Total | ~ 11 min | ~ 13 min | ~ 15 min |

### Workflow application to alternative cell types

The workflow was originally developed for *Drosophila* R7 axon terminals, which have a compact morphology. They have a single bulbous axon terminal with radially extending, relatively clearly discernible filopodial protrusion. To test the applicability of the workflow for different morpohologies, we also analyzed *Drosophila* Dm8 cells and a simulated A549-SIM cell.

#### Dm8 cell

Amacrine-like interneurons Dm8 exhibit an extended and mixed axo-dendritic branched morphology with extensive filopodial dynamics across the entire branched structure. We set the base node in the center of this branching structure and propagated it through the time steps just as in the case of R7 cells. Both cells can be seen and visually compared in Figure 9.

**Fig. 9:**
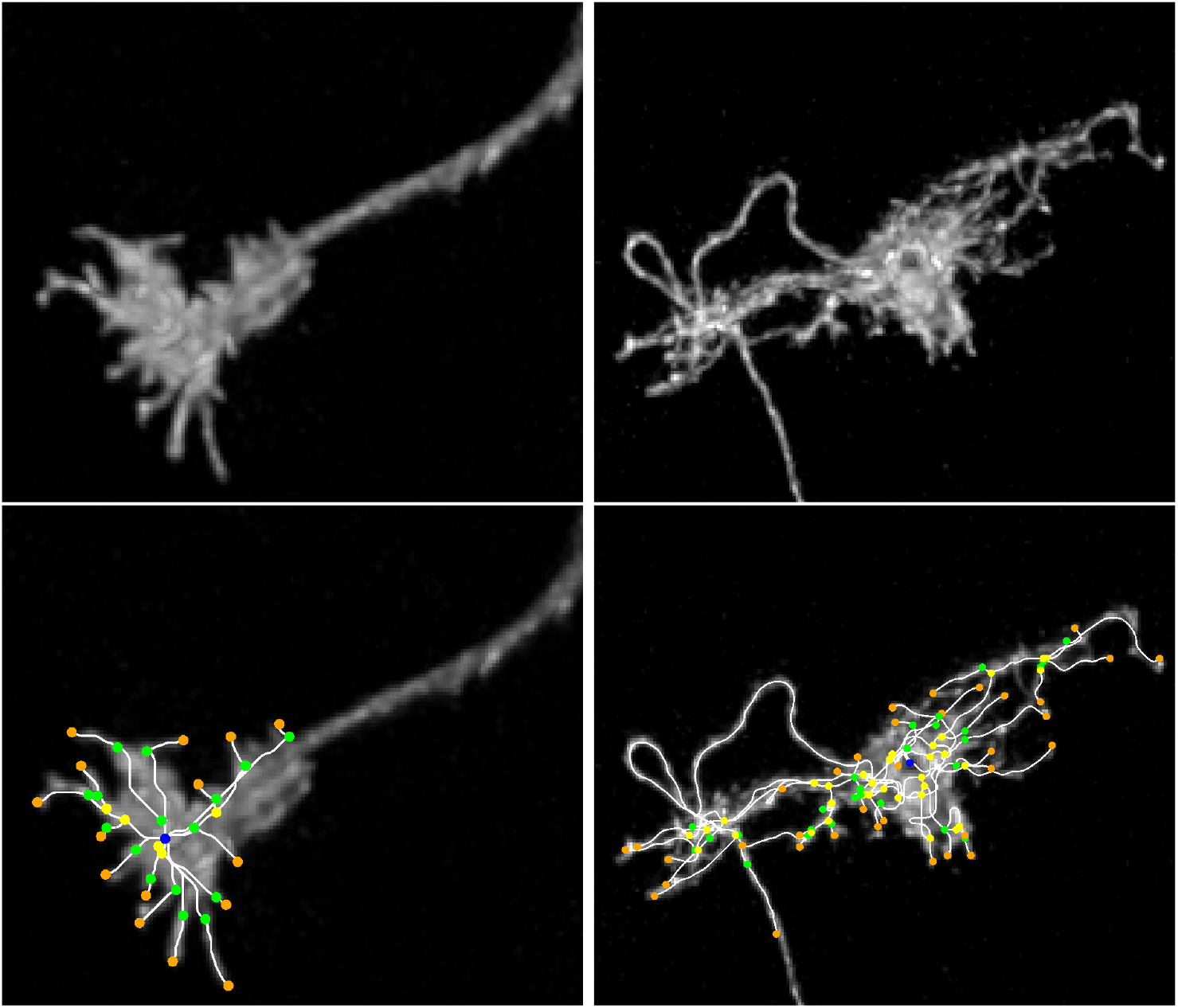
Different cell morphologies. A typical R7 cell (left) has a vastly different morphology than a typical Dm8 cell (right). Dm8 does not have an axon terminal, but an axo-dendritic branched structure which can be used for setting the root node. Reconstructions for both cell types are also shown.

The processes of tracing and tracking filopodial of Dm8s are very similar to R7 neurons. However, since Dm8 filopodia tend to be more branched than R7, they often cross and overlap. This can cause the software to trace a path from the tip to the root node that does not correspond to the traced filopodium. In such cases, it is necessary to carefully proofread the tracing results and correct potentially wrong traced paths using the implemented correction tools.

Reconstruction of the datasets was done by four neurobiologists without formal training in computational image analysis. We logged the overall time required for tracing, tracking and proofreading and used this data to analyze the required time as a function of filopodia number, lifetime and length (Fig. 10).

**Fig. 10:**
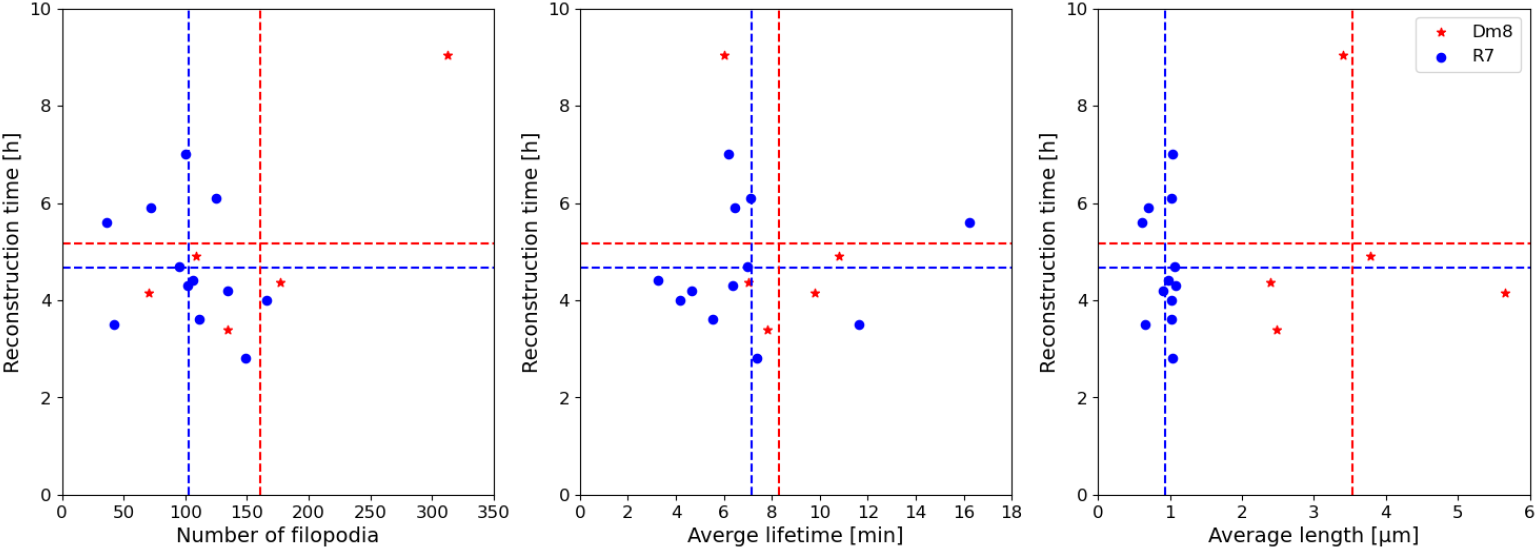
Working time comparison. To investigate the influence of filopodia properties on the time requirements, several Dm8 and R7 cells were reconstructed. The plots show reconstruction time vs. total number of filopodia, average lifetime and average length over 60 time steps. Dashed lines represent the mean values of the quantities represented by the axes.

These analyses indicate that the average working time for reconstructing a cell with 50 to 200 filopodia is approximately 3 to 5 hours, independent of the cell type. The average lifetime and length of filopodia do not correlate with reconstruction time. In a typical time step, the user works with 5 to 15 different filopodia. A video of an R7 cell, reconstructed in all time steps, is provided in the supplementary material (Video1).

#### Simulated cell

To further show that the workflow can be used for diverse cells, we have applied the workflow to the simulated cell A549-SIM from the *celltrackingchallenge*.*net*. It is important to note that the cell has a significantly different morphology and labeling compared to the R7 or Dm8 cells. The cell body is round, the inside of the cell body contains much darker values than its border, and the filopodia are extending from it linearly. Nevertheless, tracing worked robustly with the difference that traces between base nodes and root node can take unconventional paths due to the cell body being much brighter at its borders. Importantly, the tracing of filopodia worked comparably to the R7 and Dm8 neurons. Figure 11 shows a volume rendering of the cell, a cross section through its center, and the ‘filopodia’ reconstruction in a single time step.

**Fig. 11:**
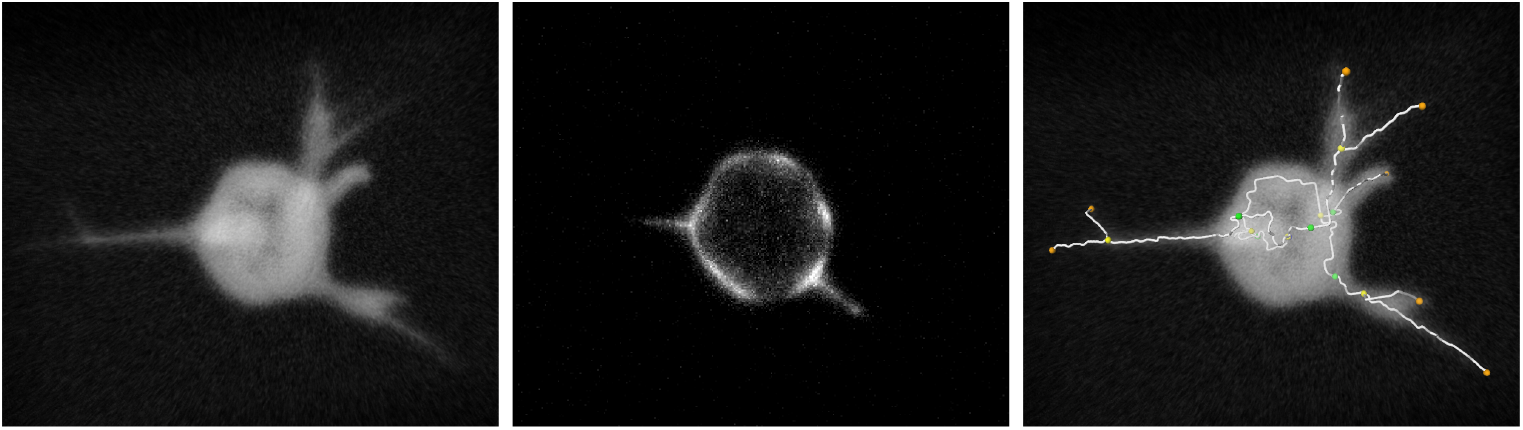
A549-SIM cell. Volume rendering of the cell (left), cross section of the cell (center), reconstruction overlaid with the cell (right).

We evaluated the reconstruction results by calculating the percentage of filopodia traces lying within the segmentation mask, which is 85% on average over all time steps. The missing 15% are all close to the base where the path is distracted due to the characteristic of the cell having bright values at the cell border but much darker values inside the cell.

## 4 Discussion

### Time requirements and applicability

The first step in the workflow is the preprocessing. The specific time needed for preprocessing depends on the dataset; in particular, the number of separate structures in a single dataset, e.g. axon terminals, results in longer manual validation times, more elaborate image cropping and Dijkstra graph calculations. The size of the axonal or branched structure also affects the preprocessing time, as larger axon structures (and hence larger cropped images) require longer calculations of the Dijkstra graphs. Both model structures analyzed here had similar preprocessing time requirements. Compared to the rest of the workflow, the preprocessing usually takes less than 5% of the total required time.

The software tool was tested by four neurobiologists who reconstructed 17 neurons of two highly divergent morphologies, namely R7 and Dm8 cells in the *Drosophila* visual system. We compared the times necessary for reconstruction (Fig. 10). On average 3 to 5 hours were necessary to reconstruct a single cell (R7 or Dm8) in 60 time steps. Neither filopodia number, lifetimes nor lengths had a significant influence on reconstruction time. The time needed for processing the automated steps is negligible compared to the time required for the manual steps.

We note that different users varied significantly in their reconstruction approaches, affecting necessary reconstruction times. To minimize bias, we applied a standardized user training protocol. However, even after training, users still varied significantly, especially with respect to the amount of time taken to validate node positions and in spotting new filopodia. Based on the difficulty of avoiding user bias, we suggest that for a given study, a single person should perform all the necessary reconstructions and thus eliminate inter-user variability. This approach is useful when relative changes between a control and experiments are more important than absolute values. When absolute values of measurements are relevant, biological criteria have to be quantitatively restricted to avoid inter-user variability.

When reconstructing a A549-SIM cell, the algorithm that generates the Dijkstra graphs needed tracing parameter adjustments to ensure the generation of satisfactory traces. We also noted that the estimation of base node locations is poorer and that they needed to be corrected more often than for the R7 or Dm8 cells. Our assumption is that the hollow body of the cell, despite the parameter adjustments, still contributes to the atypical graph, causing unconventional path generation within the cell body (as can be seen in Fig. 11). The cell body intensity distribution and the unconventional paths cause the base node to be estimated poorly. This also explains the average of 85% overlap between the traces and the segmentation mask, where the majority of the non-overlapping traces lie around the bases of the filopodia.

### Key features of the software

Our software tool was developed for tracing and tracking filopodial structures in 4D (3D+time). However, it can be applied to many different types of neuronal cells and subcellular structures without modifications. The underlying semi-automatic approach does not require training data. Apart from functioning as a stand-alone tool, the software can also serve as a means to create training data for fully automatic approaches based, for example, using deep learning [23].

Previously proposed solutions focused on 2D analysis to achieve ease of use and produce results comparably fast [17]. However, filopodia are inherently three-dimensional structures and often change growth directions in space; 2D analysis is thus likely to underestimate filopodial lengths and incorrectly represent growth directions in the observed plane. Our focus was to capture the three-dimensional movement of dynamic subcellular structures in time and provide all quantitative data of the underlying dynamics.

To facilitate the tracing and tracking in three dimensions over time, a key strength of our workflow is the ability to propagate filopodia through consecutive time steps. The user only has to validate or correct the propagated filopodia and add newly emerging filopodia when they originate. In addition, tracing of filopodial structures in space is done automatically using Dijkstra graphs; here, users only have to select the tip of a filopodium and subsequently validate the automized tracing and positioning of the base node. Results of the analysis are graphs with filopodia represented as branches originating from a root node. The representation using graphs allows practical 3D visualization and analysis of filopodia including growth direction, angle, length and branching.

All steps of the proposed workflow are supported by proofreading tools that allow the user to add, remove or modify the traced and tracked filopodia to ensure high quality output data.

### Possible improvements of the software

Validation and proofreading represent the most significant manual work and time investment using our workflow. Hence, improvements of tracing and tracking accuracy as well as facilitation of proofreading would most significantly reduce processing time. In addition, such improvement would simultaneously reduce inter-user variability.

The template matching algorithm used in the root node propagation works best for the time steps immediately following the first one. If a root node is poorly estimated for a specific time step, this error will be propagated through all the following time steps. However, for thin filopodia, the root node location does not significantly affect tracing. Future tool development may include the analysis of the dynamics of the filopodia bearing structures (in our case axon terminals or axo-dendritic branched structures). Possible implementations could include elastic image registration or the use of machine learning.

Finally, significant workflow improvements are possible by improving the interface based on feedback from experienced users. For example, a more interactive way of entering corrections of filopodial nodes or tracks using a simple “drag-and-drop” approach could replace the manual selection of an object and moving it to a new location through the graphical interface.

### Tool’s significance for cellular and developmental neuroscience

The development and application of advanced 4D live image acquisition in neuroscience, as elsewhere, has increased substantially in recent years. New microscopy and non-invasive imaging technologies allow to obtain live imaging data of three-dimensional structures with increasing temporal resolution. However, 4D data analysis and interpretation have remained far behind these developments. For example, tracking of subcellular structures in three-dimensional data over time has remained a largely unmet challenge and currently represents a substantial barrier to progress [32]. Similarly, the extraction of principles from 4D data typically requires quantitative data analysis for subsequent modeling.

All across cell biology, dynamic processes in three dimensions underlie mechanisms that may not be fully understood based on fixed data. Neurons in particular are highly polarized cells with long-range trafficking and morphogenetic dynamics that play critical roles during development and function. In this work, we focused on the dynamics of axonal growth cones and axo-dendritic arborizations, structures whose dynamic properties may directly influence the development of brain wiring [1]. For example, the development of brain wiring is still mostly studied based on adult outcomes, rather than during the dynamic period, before, during and after synaptic partner choice, which is certainly, at least in part, due to the technical challenges involved [33].

## 5 Conclusion

We developed a semi-automatic workflow for the time-efficient tracing and tracking of filopodia in 4D microscopy data (3D+time). In this workflow, most of the tasks are automized and the majority of the user input is restricted to validating parameters and results. The implemented software tool supports this proofreading by providing several options for interactive corrections. Finally, filopodial structures, like the ones at the center of this study, are not restricted to neurons. Many motile cell types utilize filopodial dynamics to explore and move in their environments. While many in vitro studies focused on cellular movement in artificial two-dimensional environments, the migration and underlying dynamics of motile cells are now increasingly accessible in the three-dimensional organismal context. The tool developed here was devised to facilitate such a diverse range of applications. Its application to the R7, Dm8 and A549-SIM datasets showed robust function and wide ranging applicability for filopodial tracing across different datasets. The tool is publicly available at https://github.com/zibamira/filopodia-tool along with the user manual and example datasets.

## Supporting information

A video of an R7 cell, reconstructed in all time steps, is provided in the supplementary material (Video1).

## 6 Availability and requirements

**Project name:** 3D filopodia tracing and tracking tool

**Project home page:** https://github.com/zibamira/filopodia-tool

**Operating system(s):** Platform independent

**Programming language:** C++

**Other requirements:** Development version of Amira software

**License:** MIT-style

**Any restrictions to use by non-academics:** None

## 7 List of abbreviations

2D: Two dimensions or two-dimensional
3D: Three dimensions or three-dimensional
4D: Four dimensions or four-dimensional
CTS: Contour Tree Segmentation
NCC: Normalized Cross Correlation
RMSD: Root Mean Squared Deviation

## 8 Declarations

### Ethics approval and consent to participate

Not applicable.

### Consent for publication

Not applicable.

### Availability of data and materials

Two example datasets are provided in the online repository (https://github.com/zibamira/filopodia-tool). Additional datasets and results can be provided upon request.

### Authors’ contributions

BB: Methodology, software, formal analysis, writing (original draft, review & editing)

JB: Methodology, software, formal analysis, writing (original draft, review & editing)

VJD: Software, conceptualization, supervision, writing (review & editing)

MNÖ : Data curation, methodology, investigation, writing (review & editing)

AK: Data curation, methodology, investigation, writing (review & editing)

NW: Data curation, methodology, investigation, writing (review & editing)

SP: Conceptualization, funding acquisition, project administration, supervision, writing (review & editing)

PRH: Conceptualization, funding acquisition, project administration, writing (review & editing)

DB: Methodology, software, formal analysis, conceptualization, funding acquisition, supervision, writing (review & editing)

## Acknowledgements

We would like to thank all members of the Baum and Hiesinger labs as well as members of the Research Consortium *RobustCircuit* for discussion. We are particularly grateful to Joachim Fuchs for providing feedback to the manuscript and Justus Vogel for supporting the development of the Amira software.

